# Evolutionary Origins of Recurrent Pancreatic Cancer

**DOI:** 10.1101/811133

**Authors:** Hitomi Sakamoto, Marc A. Attiyeh, Jeffrey M. Gerold, Alvin P. Makohon-Moore, Akimasa Hayashi, Jungeui Hong, Rajya Kappagantula, Lance Zhang, Jerry Melchor, Johannes G. Reiter, Alexander Heyde, Craig M. Bielski, Alexander Penson, Debyani Chakravarty, Eileen M. O’Reilly, Laura D. Wood, Ralph H. Hruban, Martin A. Nowak, Nicholas D. Socci, Barry S. Taylor, Christine A. Iacobuzio-Donahue

**Affiliations:** Sloan Kettering Institute, Memorial Sloan Kettering Cancer Center, NY NY USA; Department of Pathology, Memorial Sloan Kettering Cancer Center, NY NY USA; Department of Surgery, Memorial Sloan Kettering Cancer Center, NY NY USA; Department of Medicine, Memorial Sloan Kettering Cancer Center, NY NY USA; Department of Epidemiology and Biostatistics, Memorial Sloan Kettering Cancer Center, NY NY USA; Bioinformatics Core, Memorial Sloan Kettering Cancer Center, NY NY USA; Human Oncology and Pathogenesis Program, Memorial Sloan Kettering Cancer Center, NY NY USA; David M. Rubenstein Center for Pancreatic Cancer Research, Memorial Sloan Kettering Cancer Center, NY NY USA; Marie-Josee and Henry R. Kravis Center for Molecular Oncology, Memorial Sloan Kettering Cancer Center, New York, NY USA; Program for Evolutionary Dynamics, Harvard University, Cambridge MA USA; Department of Mathematics and Department of Organismic and Evolutionary Biology, Harvard University, Cambridge MA USA; Canary Center for Cancer Early Detection, Department of Radiology, Stanford University, Palo Alto CA USA; Department of Pathology, Johns Hopkins Medical Institutions, Baltimore MD USA; Sol Goldman Pancreatic Cancer Research Center, Baltimore MD USA

## Abstract

Surgery is the only curative option for Stage I/II pancreatic cancer, nonetheless most patients will recur after surgery and die of their disease. To identify novel opportunities for management of recurrent pancreatic cancer we performed whole exome or targeted sequencing of 10 resected primary cancers and matched intrapancreatic recurrences or distant metastases. We identified that adjuvant or first-line platinum therapy corresponds to an increased mutational burden of recurrent disease. Recurrent disease is enriched for mutations that activate Mapk/Erk and PI3K/AKT signaling and develops from a monophyletic or polyphyletic origin. Treatment induced genetic bottlenecks lead to a modified genetic landscape and subclonal heterogeneity for driver gene alterations in part due to intermetastatic seeding. In one patient what was believed to be recurrent disease was an independent (second) primary tumor. These findings advocate for combination therapies with immunotherapy and routine post-treatment sampling as a component of management of recurrent pancreatic cancer.

## Introduction

Pancreatic ductal adenocarcinoma (PDA) is currently the 3th leading cause of cancer death in the United States and is projected to become the 2nd cause of cancer death within five years (1,2). Several reasons account for these statistics, including an inability to diagnose the disease when at a curative stage, late presentation and modest impact of current best available therapies (3). There is a limited understanding of the genetics of recurrent disease which limits targeted therapy opportunities or informed design of clinical trials.

Approximately 10-15% of newly diagnosed PDA patients are diagnosed with early-stage disease (Stage I or II). For these patients, surgical resection followed by adjuvant therapy is the only option for cure (3). While long term survival following resection of PDA has been reported (4–6), the majority of patients who undergo resection will recur locally or at distant sites and die of their disease within five years. Several factors have been shown to have predictive or prognostic value for disease free or overall survival in resected PDA patients, including a high ratio of involved to total resected lymph nodes, larger tumor size, high tumor grade, the presence of vascular and perineural invasion, or variably positive margins (7–9). Venous invasion is very common in pancreatic cancer and may contribute to the aggressiveness of this disease (10). Molecular features of PDA have also been attributed to worse outcome after surgery. For example, patients with co-incident *TP53* and *SMAD4* alterations have shorter disease-free survival than patients whose tumors do not have these genetic alterations (11–13). Alternatively, the presence of a basal expression signature (14–16), paucity of an immune signature (17) or microbial dysbiosis (18) have also been associated with worse overall survival.

Ultimately the extent to which recurrent PDA has genetic features of clinical significance or potential actionability is unknown. To address this question and to improve our understanding of recurrent PDA we performed whole exome and/or targeted sequencing of 10 primary PDAs, matched local (pancreatic resection bed) recurrences and multiple anatomically diverse metastases. We identified that pancreatic cancer recurrences following surgery have an increased mutational burden, distinct subclonal origins and in some instances are characterized by somatic mutations with potential implications for clinical management.

## Results

We screened a collection of more than 160 PDA research autopsies to identify patients for whom a sample of their original surgical pancreatic resection was available, who underwent adjuvant treatment after surgery, and who had histologically confirmed recurrent disease within the pancreatic remnant and one or more metastases to anatomically distinct sites such as the liver, lungs or peritoneum (Fig. 1A,B). We identified nine such patients for study (Table S1). An additional patient was included that did not have metastatic disease at autopsy but did have an aggressive local recurrence with multiple geographically distinct samples of this mass available for profiling. One normal tissue sample from each patient was also used to distinguish somatic from germline variants (Table S2).

**Figure 1.**
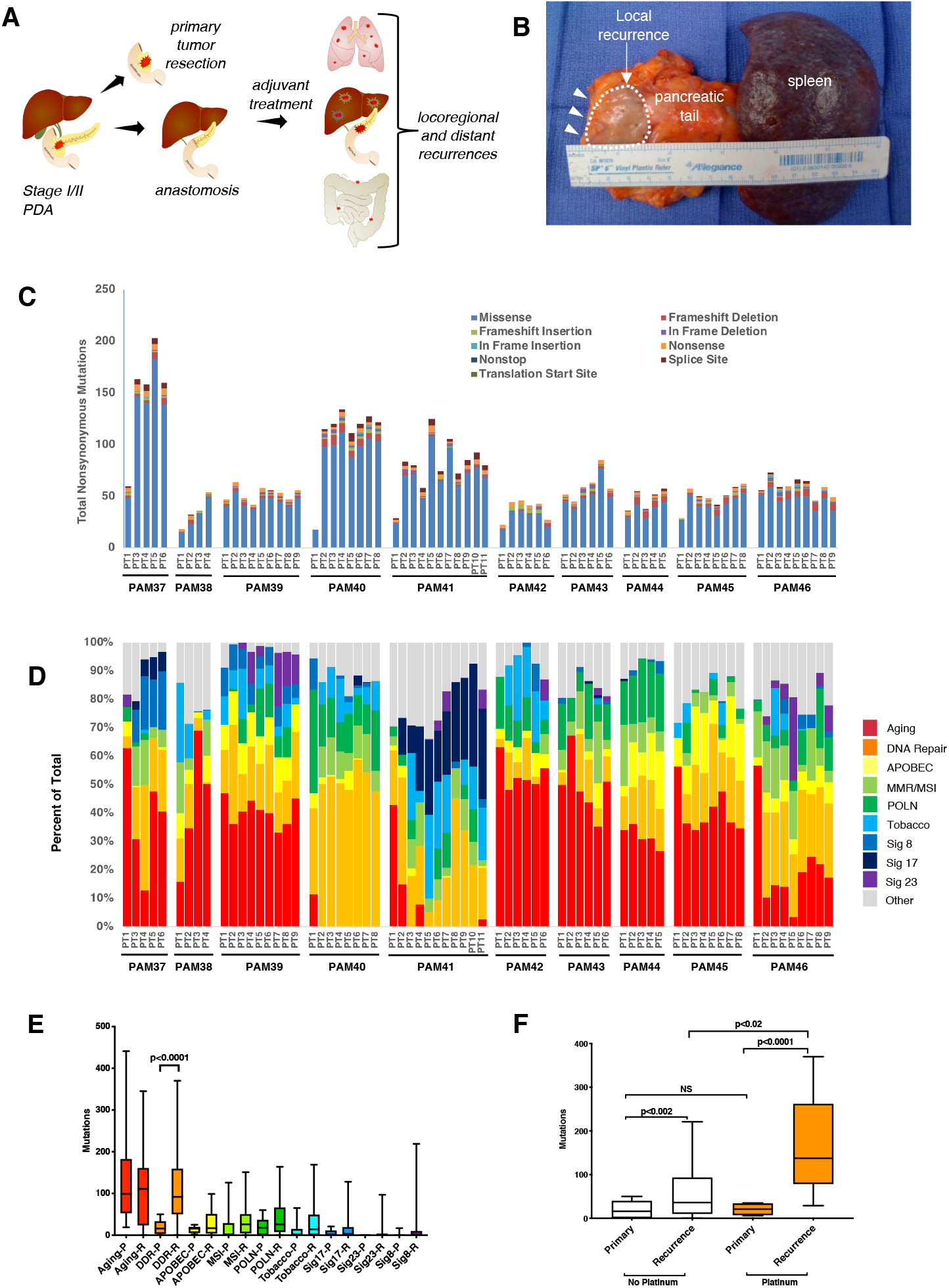
Recurrent pancreatic cancer after adjuvant therapy has distinct genetic features compared to the primary tumor. **A.** Schematic illustrating the temporal and spatially distinct sample types collected for this study. **B.** Pancreatic tail remnant and attached spleen removed at autopsy of patient PAM41. Arrowheads indicate the resection margin and dashed circle indicate the location and dimensions of the recurrent tumor. **C.** The number and type of nonsynonymous somatic alterations identified in each sample for each patient. **D.** Most common mutational signatures identified across the entire sample cohort. **E.** Comparison of the prevalence of distinct mutational signatures in primary carcinomas versus recurrent disease indicates a statistically significant difference in distribution between the two groups (two-sided χ^2^-squared test, p<0.0001). A statistically significant increase in mutations characteristic of the DNA damage repair signature are also present in recurrent disease (p<0.0001, Mann-Whitney U test_ **F.** Comparison of DNA damage repair signature in patients who did and did not receive a platinum agent as adjuvant or first line therapy. All comparisons by Mann-Whitney U test.

All samples were analyzed by whole exome sequencing (WES) to a median of 330x coverage (range 135-652x)(see Methods, Table S3). To supplement the breadth of WES sequencing in these samples, we also performed concurrent deep sequencing using a targeted sequencing panel to ensure greater sensitivity for mutations and DNA copy number alterations in 410 cancer associated genes (Table S4) (19). We identified 4864 total somatic single nucleotide variants (SNVs) and small insertions or deletions (indels) with a median of 57 per sample (range 17-203; Fig. 1C, Table S5). We found no evidence of microsatellite instability in these 10 patients, nor were germline or somatic mutations identified in recognized MSI-related or other DNA damage repair genes (20).

To better understand the patterns of mutation accumulation in primary versus recurrent disease, we performed mutational signatures analysis of all samples that underwent WES (Fig. 1D). Nine signature classes were identified in our cohort: aging (signatures 1 and 5), double strand break repair (DSBR, signature 3), apolipoprotein B mRNA editing enzyme, catalytic polypeptide-like (APOBEC, signatures 2 and 13), mismatch repair defects (signatures 6, 15, 20, 21 and 26), tobacco (signature 4), somatic hypermutation (signature 9), and the three unknown signatures 8, 17 and 23 (21). A high prevalence of Signature 9 (POLN) was seen in at least one sample in seven of eight patients. This signature is associated with somatic hypermutation by polymerase η, a DNA polymerase that plays a role in DNA repair by translesion synthesis (22). As previously reported, we also identified subsets of patients with an abundance of signatures with unknown etiology such as signatures 8 and 17 (17). One patient (PAM41) had a remarkable abundance of mutations characteristic of signature 23, a rare type of signature of unknown etiology (21). A comparison of the prevalence of each of these nine signatures in primaries to recurrent tumors revealed that a statistically different distribution in treated disease (two-sided χ^2^-squared test, p<0.0001, Fig. 1E). Comparisons of the prevalence of each signature class specifically in post-treatment recurrent disease versus treatment naïve primary carcinomas revealed a significant increase in the double strand break repair signature (median 16 somatic mutations per primary versus 92 in recurrent disease, Mann-Whitney U test, p<0.0001). Four patients received a platinum agent as part of their adjuvant or first line therapy for recurrent disease (Table S1), thus we determined the extent that this signature was enriched in these patients compared to those who received other regimens. Comparison of these two groups revealed that patients who did not receive a platinum agent had a modest yet significant increase in their recurrent disease with respect to the double strand break repair signature (median 58 somatic mutations per primary versus 532 in recurrent disease, Mann-Whitney U test, p<0.002), and this signature was remarkably more prevalent in the recurrent disease of patients who received one or more platinum agents (median 84.5 somatic mutations per primary versus 1379 in recurrent disease, Mann-Whitney U test, p<0.0001) (Fig. 1F).

We next determined the somatic alterations in known cancer genes in each patient. We identified somatic mutations in known PDA driver genes predicted to have functionally deleterious effects such as those in *KRAS*, *CDKN2A*, *TP53*, *SMAD4*, and *ARID1B* (Table S6 and S7). Collectively, the genetic features of the resected PDA samples (sample PT1 from each patient) were consistent with previous sequencing studies of these cancers (15). By contrast, the genomic features of recurrent disease (Fig. 2, Table S6) were notable for somatic alterations in additional genes, some of which are predicted to or likely to activate Mapk/Erk signaling (G12V mutation in *KRAS,* I679Dfs*21 mutation in *NF1*, R111X mutation in *PPP6C*)(23,24), PI3K/Akt/MTOR signaling (D323H hotspot mutation in *AKT1*, E542K hotspot mutation in *PIK3CA*, homozygous deletion of *STK11*)(25), or MYC/MAX regulated gene expression (Y1312X mutation in *CHD8,* W1004X mutation in *MGA, X863_* splicing mutation in *NOTCH1*, up to 16-fold amplification of *MY*C) (26,27). Somatic alterations in genes predicted to affect chromatin-mediated gene expression (up to 38-fold focal amplification of *HIST1H3B*, frameshift mutations in *KMT2B*, *KMT2C, KMT2D,* c.4471-1N>A splicing mutations in *TRIP12*) (28–30), nuclear export (E571K hotspot mutation in *XPO1)* (31) and DNA damage repair (c.497-1N>A splicing mutation in *ATM,* F134V mutation in *TP53*) were also found (32,33). Recurrences from two patients contained copy number alterations of genes implicated in innate immunity signaling (*PRKCI* 38-fold focal amplification, *TRAF3* homozygous deletion) (34,35). We also noted that whole genome duplication (WGD) was present in one or more samples of recurrent disease in eight patients (36). In only two of these eight patients was WGD detected in the primary tumor indicating gains in ploidy may be common during PDA progression. Collectively these data identify potential signaling and transcriptional pathways by which recurrent disease develops; with some exceptions these alterations are mutually exclusive across PDAs.

**Fig. 2.**
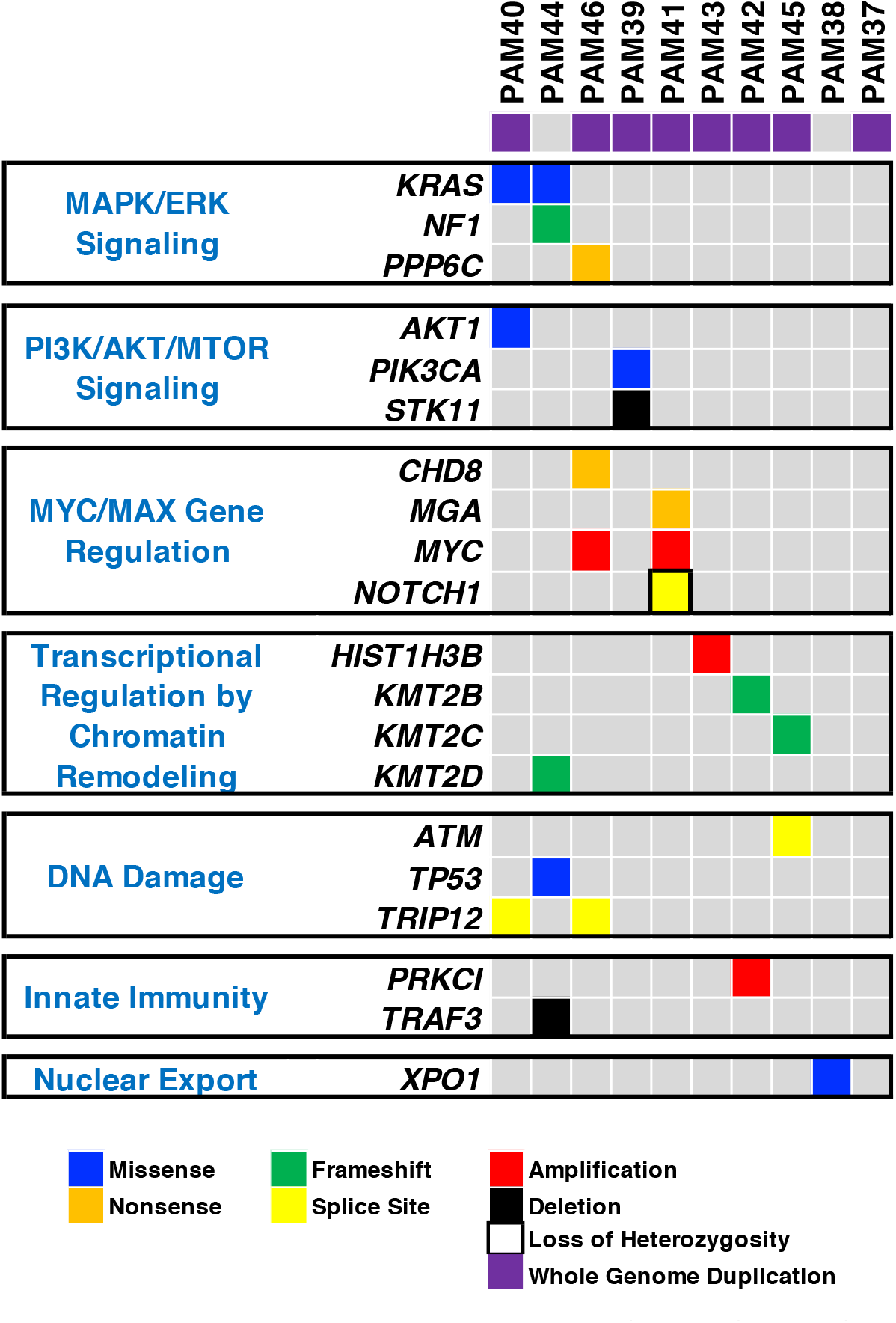
Oncoprint of Recurrent Pancreatic Cancer. Somatic mutations and copy number alterations present in one or more samples of the recurrent PDA sampled at autopsy but not in the matched originally resected primary tumor. Somatic alterations present in both the primary and recurrent disease are listed in Tables S6 and S7.

We hypothesized that somatic alterations identified in recurrent PDA samples reflect the clonal expansion of pre-existent populations following selective pressure imposed by adjuvant therapy. Pairwise cancer cell fraction (CCF) plots generated for all patients confirmed the enrichment of one or more subclonal populations in the recurrent disease compared to the primary tumor (Fig. 3A). We specifically focused on the *AKT1* D323H in PAM40, *PIK3CA* E542K in PAM39 and *NOTCH1* X863_splice site mutation in PAM41 as they represent functionally relevant alterations and in theory may be clinically actionable (http://oncokb.org). The *KRAS* p.G12V mutation in PAM40 was also of interest given recent reports of subclonal *KRAS* mutations in PDA (15). Droplet digital PCR confirmed that in all instances these mutations existed in the primary tumor at prevalence rates from 0.2-2%, below the level of detection of our WES (Fig.3B-D).

**Figure 3:**
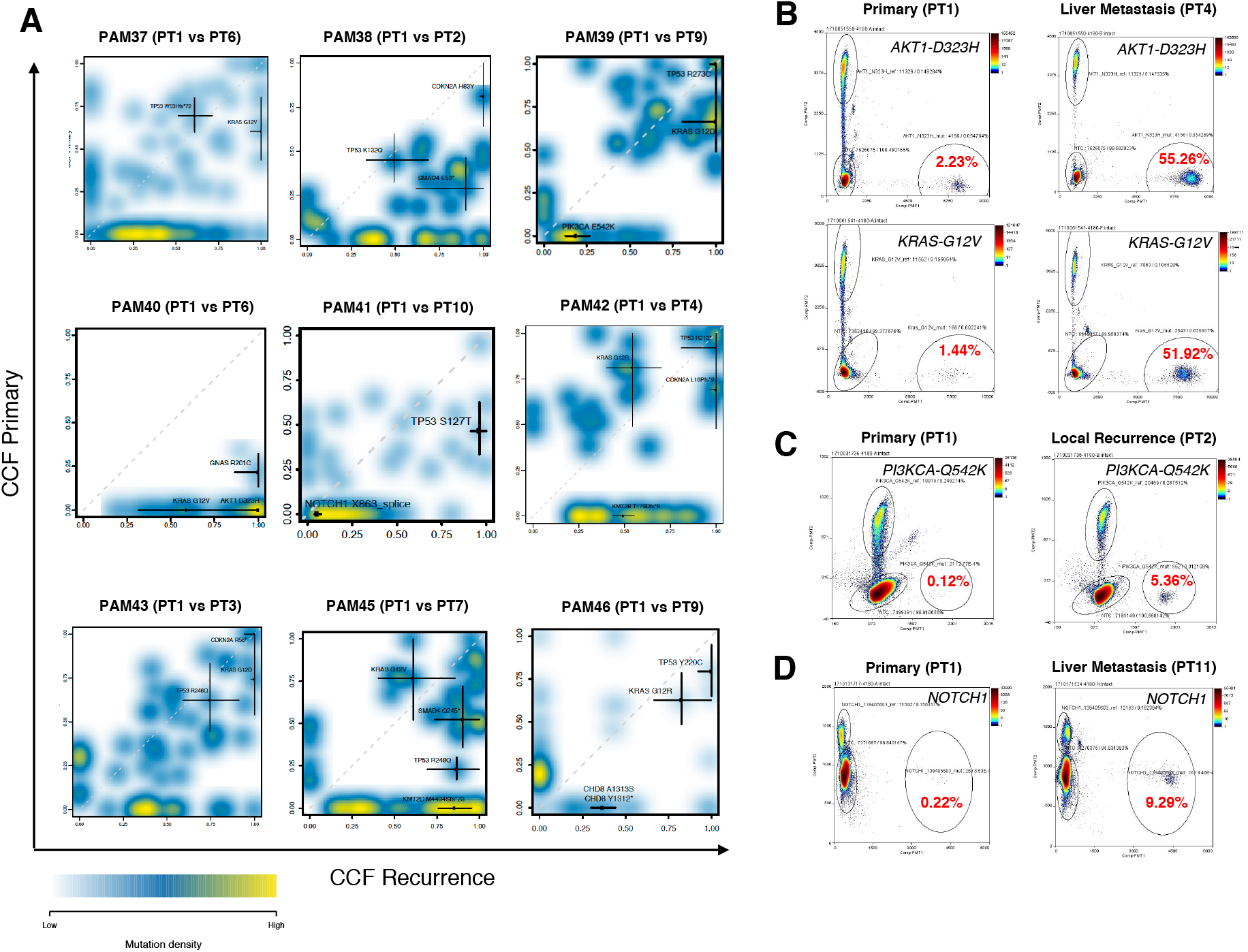
Somatic alterations in recurrent pancreatic cancer alterations reflect expansion of subclones pre-existent in the primary tumor. Representative density cloud plots of the primary tumor (PT1) and one matched sample of recurrent disease for nine different patients. Subclonal expansion of cells containing a *PIK3CA* E524K mutation (PAM39), a *KRAS* G12V and *AKT* D323H mutation (PAM40), a *NOTCH*1 frameshift mutation (PAM41), *KMT2C* frameshift mutation (PAM45) and *CHD8* nonsense and missense mutations (PAM46) in the recurrent disease are seen. The cancer cell fractions of representative truncal driver genes for all cases (i.e. KRAS, TP53, SMAD4 and/or GNAS) are shown for reference. reference. **B-D.** Droplet digital PCR analysis of mutant allele abundance in the primary tumor and one matched sample of recurrent disease in three different patients. In all three patients subclones were preexistent at low allele frequencies in the primary tumor. Each dot represents one droplet, and color bars at top right of each plot indicate relative intensity of the VIC labeled mutant (x axis) and FAM labeled wild type allele (y axis) fluorescent labels.

To understand the evolutionary origins of recurrent disease in each patient we inferred phylogenies based on high confidence mutations in each patient (Methods)(37). One patient who was clinically thought to have an intrapancreatic recurrence of the resected PDA after 18 months, but molecular analysis revealed a second (independent) primary PDA (Fig. 4A). For example, the resected primary tumor (PAM44PT1) had a *KRAS* p.G12R mutation and 80 private passenger mutations, whereas the “recurrence” samples from this patient (PAM44PT2-PT5) shared 124 mutations not seen in the primary. These latter mutations included a *KRAS* p.G12D, a *TP53* p.F134V and a 29 basepair frameshift deletion in *KMT2D* (Fig. 4B). A CCF plot of the original primary (PAM44PT1) compared to the “recurrence” samples confirmed the mutual exclusivity of the somatic mutations in each lineage (Fig. 4C). Re-review of the histology of the first primary tumor and associated imaging studies did not suggest the presence of a cystic neoplasm. These findings are highly indicative of two independent primary tumors that arose from distinct precursors (38).

**Figure 4:**
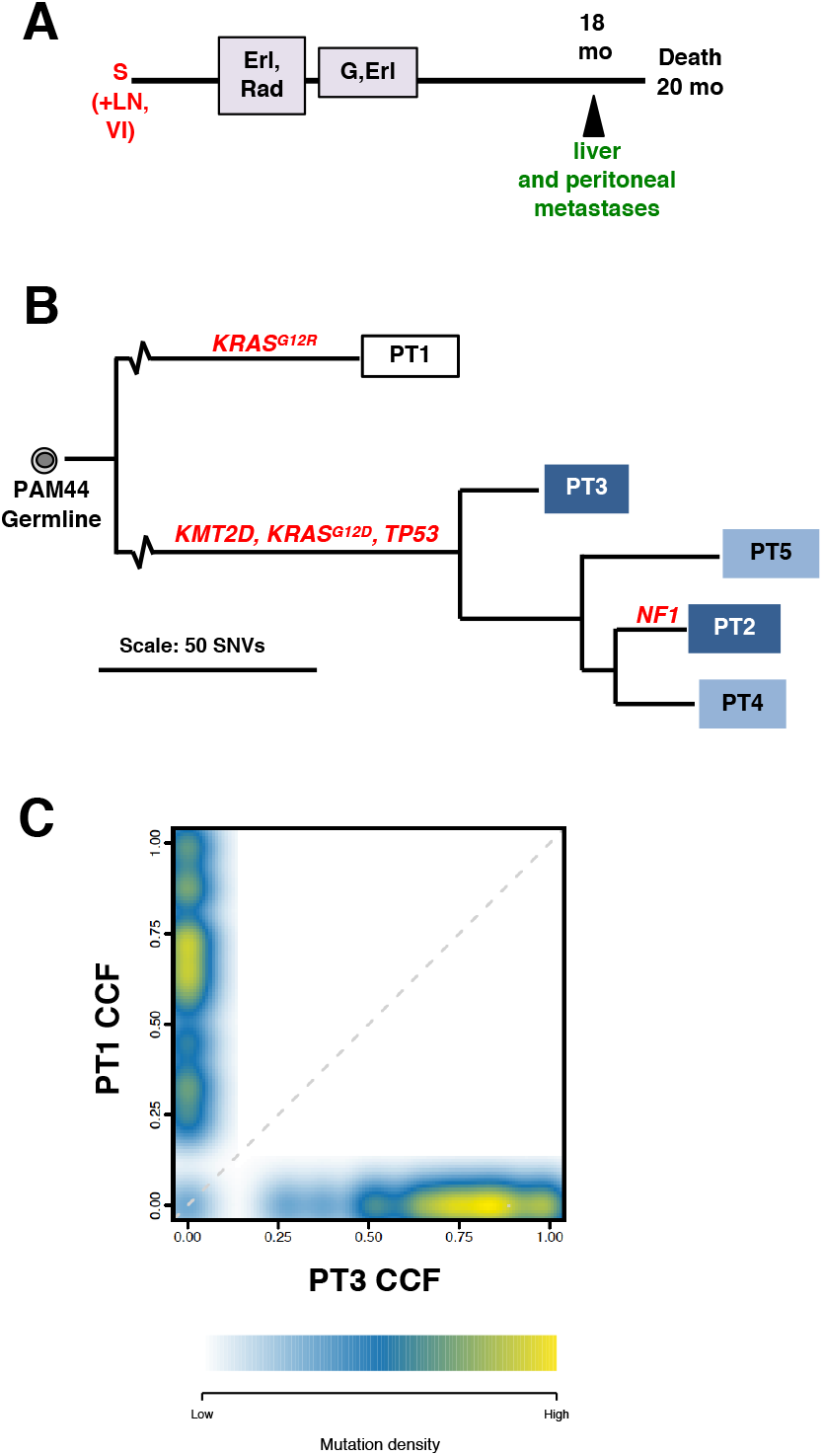
Metachronous pancreatic cancer simulating a local recurrence. **A.** Clinical timeline of events in this patient. Liver and peritoneal metastases were radiographically evident at 18 months and a mass in the pancreatic remnant was noted at autopsy. **B.** Phylogenetic analysis of whole exome sequencing data generated for this patient indicates the metachronous pancreatic cancer contains a unique repertoire of somatic driver gene alterations including a *KRAS-G12D* mutation compared to the first primary that contains a G12V mutation. **C.** Representative density cloud plot of the first primary tumor (PT1) and one sample of the metachronous primary tumor (PT3). The majority of somatic alterations in each primary are mutually exclusive from each other.

In the remaining nine patients the sample of resected primary tumor and all samples of recurrent disease arose from a common ancestor of neoplastic cells containing canonical PDA driver mutations. However, there were two distinct evolutionary trajectories by which recurrent disease arose. For five patients (PAM37, PAM38, PAM40, PAM42, PAM46) the resected primary tumor sample formed the outgroup in the phylogeny (Fig.5A,B and Figs. S1) indicating that in these patients the recurrent disease developed from a single residual clonal population, i.e. a monophyletic origin. In the remaining four patients (PAM39, PAM41, PAM43, PAM45) (Fig. 5C,D and Fig. S2) the recurrent disease was inferred to be seeded by multiple ancestral clones and was polyphyletic in origin. For a more objective metric of relatedness among the primary and recurrent disease in each patient we calculated pairwise Jaccard similarity coefficients for all samples within a patient. The average Jaccard indices per patient supported the distinction of monophyletic versus polyphyletic origins of recurrent disease (Fig. 5E-G) in that monophyletic recurrences were significantly more distant (divergent) from the primary tumor, whereas polyphyletic recurrences were highly related to the primary tumors. Recognizing the sample numbers are low for robust outcomes analysis, exploratory analysis indicates that patients with monophyletic recurrences had a longer disease-free but not overall survival (Fig. 5H,I). From phylogenetic analysis alone, the timing of dissemination to other organs cannot be readily resolved. However, utilizing mathematical modeling and previously measured metastasis doubling times (39), we found that the minimal time required to grow from one to a billion cells (roughly 1 cm^3^) is 1.82 years, or 21.9 months (90% CI: 1.61 – 2.05 years) (Methods). Since clinical metastases occurred much earlier than these required 1.82 years after surgery in all patients with distant disease (median metastasis free survival 11.0 months, range 6-18 months)(Table S1), at least one of the metastases must have been microscopically seeded before surgery consistent with prior estimates (40). Patient PAM46 typified these dynamics as they had a grossly positive surgical margin and developed a radiographically evident locoregional recurrence, but no metastases, 17 months after surgery (Fig. S1J-L). Irrespective of origin, in all nine patients additional subclonal expansions occurred after dissemination, in some cases resulting in spatial heterogeneity for driver gene mutations among the different sites of recurrent disease (Fig. 5 and Figs. S1,S2).

**Figure 5:**
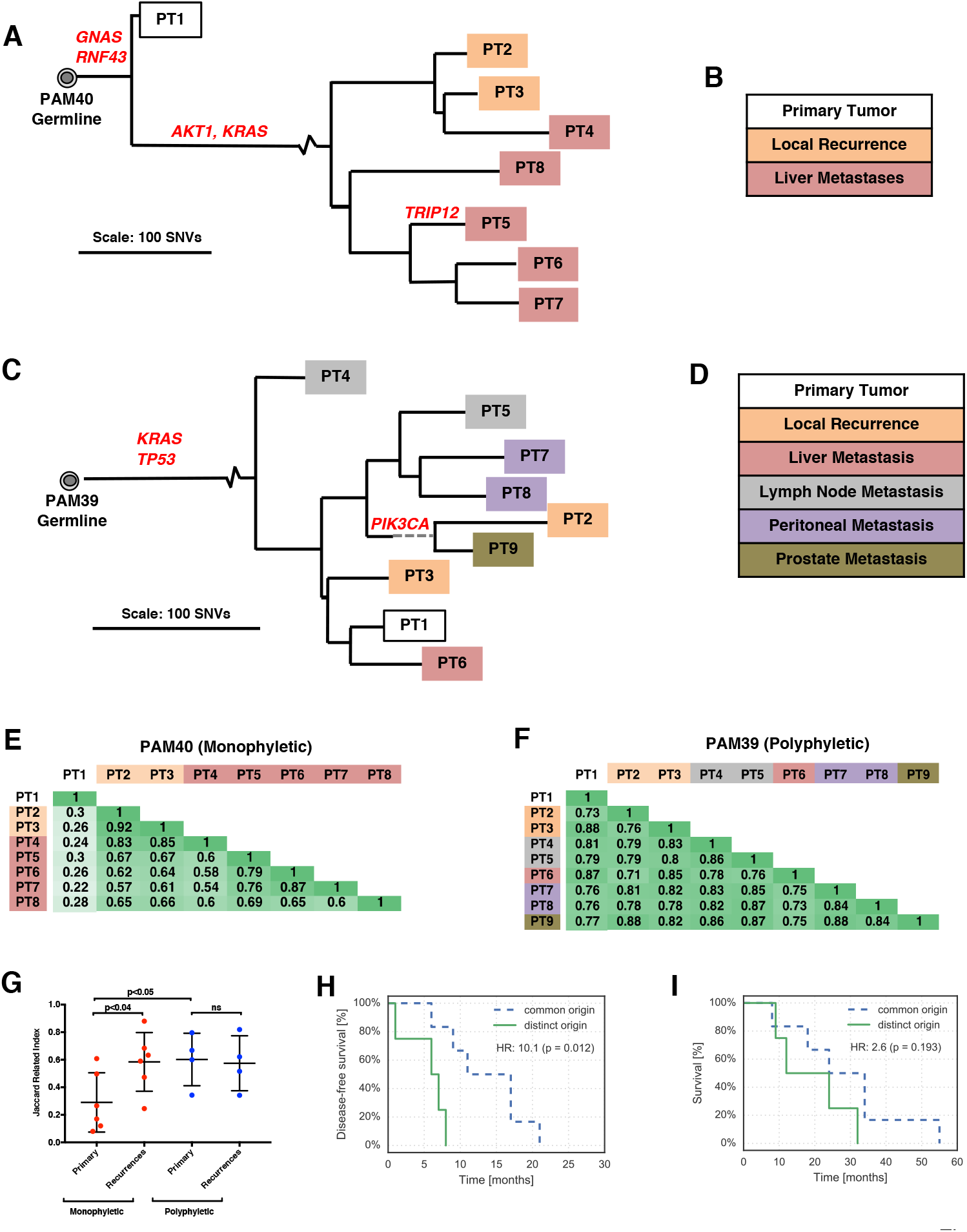
Recurrent pancreatic cancer originates from two distinct evolutionary origins. **A.** Phylogenetic analysis of the relationships of the primary tumor to the local recurrence and liver metastases in patient PAM40. All recurrent disease is the result of clonal expansion of a single pre-existent subclone notable for an *AKT1* and *KRAS* mutations (See Fig 2B). The primary tumor (PT1) is the outgroup in the tree. A subclonal *TRIP12* mutation is also seen in a single liver metastasis. **B.** Color code of sample origins in PAM40. **C.** Phylogenetic analysis of the relationships of the primary tumor to the local recurrence and liver metastases in patient PAM39. In this patient the recurrent disease is the result of more than one clonal expansion. The preexistent *PIK3CA* mutation has expanded in the lineage that gave rise to samples PT2 and PT9. **D.** Color code of sample origins in PAM39. **E,F.** Jaccard indices for each pairwise comparison in PAM40 (monophyletic recurrence) versus PAM39 (polyphyletic recurrence). **G.** Comparison of the average Jaccard index for primary tumors and their matched recurrences in patients with monophyletic recurrences versus those with polyphyletic recurrences. Monophyletic recurrences are significantly different from their matched primary tumor indicating passage through a genetic bottleneck, whereas no difference is found between the primary tumor and matched recurrences in patients with polyphyletic recurrences. (comparisons by a Student’s two-sided T test). Monophyletic (“common origin”) recurrences are associated with an improved disease-free survival (**H**) but not overall survival (**I**) in this small cohort. (comparisons by log-rank test of Kaplan Meier survival curves).

We next sought to understand the clonal origins of local recurrences, a major clinical issue for patients who undergo potentially curative resection (41). We therefore inferred the migration patterns of recurring disease across spatially distinct sites in each patient (Methods). These analyses indicated that the seeding patterns of recurrent disease were diverse, with potentially multiple patterns arising even in the same patient. Metastasis to metastasis seeding was evident in some patients, typified by PAM37 in whom an omental metastasis seeded three liver metastases (Fig. 6A), in PAM38 in whom a liver metastasis seeded a lung metastasis (Fig. 6B), and in PAM45 in whom a perirectal metastasis seeded two abdominal wall metastases (Fig 6C). Local recurrences can also be seeded by the primary tumor or by a metastasis. In PAM41 (Fig. 1B and Fig. 7A) and PAM43 (Fig. 7B), the primary tumor seeded the local recurrence despite both having negative surgical margins at the time of resection. By contrast, PAM38 (Fig. 6B) and PAM42 (Fig. 7C) had local recurrences seeded by a liver or lung metastasis, respectively. Analysis of PAM39 and PAM40 indicated that two or more seeding patterns were equally likely to have occurred for all sites analyzed and thus no dissemination events could be confidently inferred. Inter-metastatic seeding was inferred in patients with both monophyletic (PAM37, PAM38, PAM42) and polyphyletic (PAM41, PAM43, PAM45) recurrences collectively indicating that migration patterns are unrelated to phylecity of the recurrent disease. While we studied a small cohort, surgical margin status appeared to be unrelated to the origin of a local recurrence. For example, local recurrences seeded by the primary tumor were noted in patients with negative surgical margins (i.e. PAM41, Fig. 7A) whereas local recurrences seeded by a metastasis were noted in patients with a positive surgical margin (i.e PAM38, Fig. 6B).

**Figure 6:**
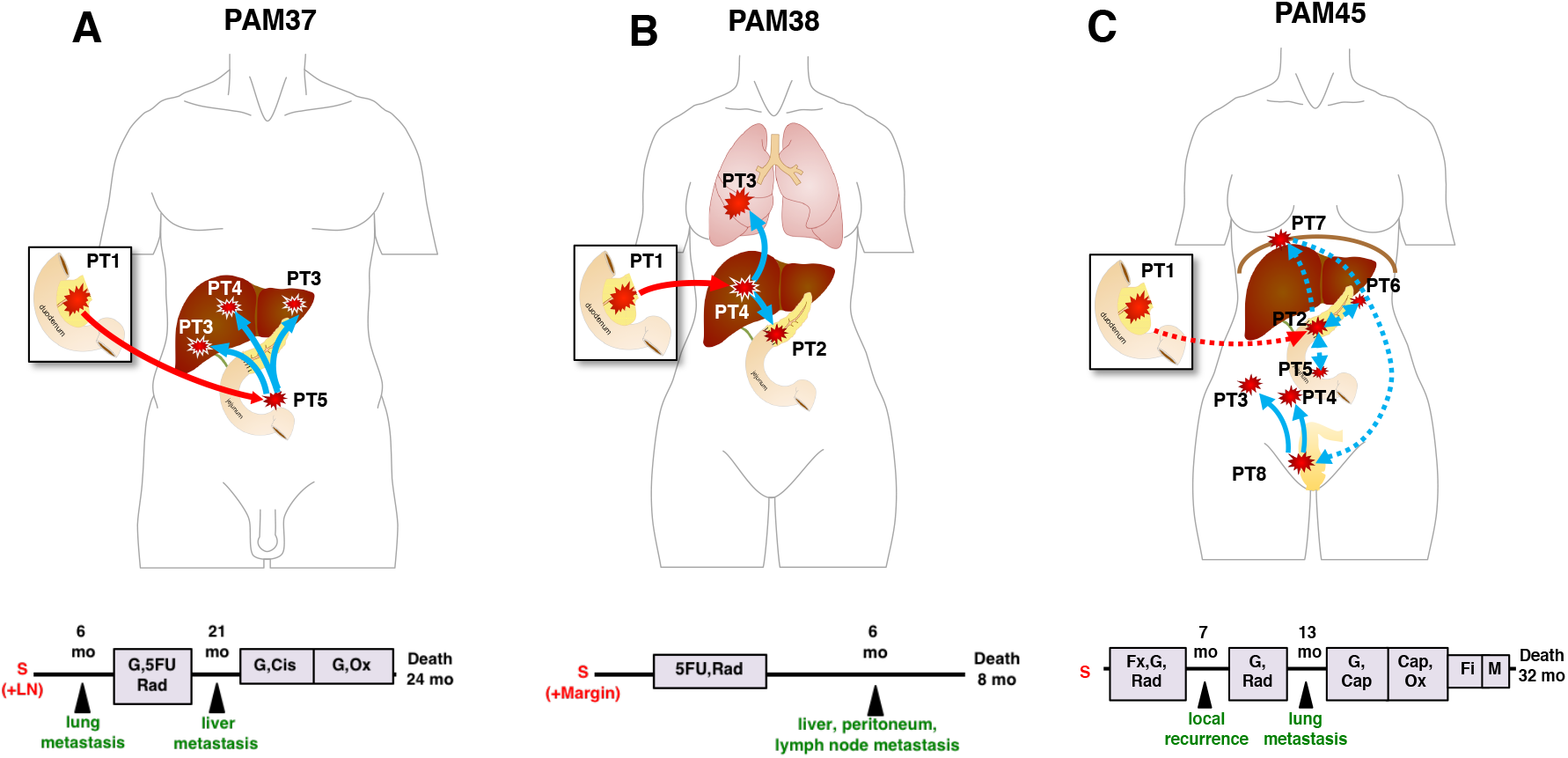
Intermetastatic seeding in recurrent pancreatic cancer in patients PAM37 (A), PAM38 (B) and PAM45 (C). The previously resected primary tumor is indicated by PT1 to the left of each patient schematic, and each patients’ clinical course is indicated below. Abbreviations are G, gemcitabine; Rad, radiation; Fx, FOLFIRINOX, Fi, FOLFIRI; Cis, cisplatin; Cap, capecitabine; Ox, oxaliplatin; M, Mek Inhibitor. Solid lines indicate high confidence migration patterns and dashed lines indicate low confidence migration patterns inferred by MACHINA. Red lines reflect migration from the primary tumor and blue lines indicate migration from a site of recurrent disease. Each patient had at least one high confidence migration event from one site of recurrent disease to another.

**Figure 7:**
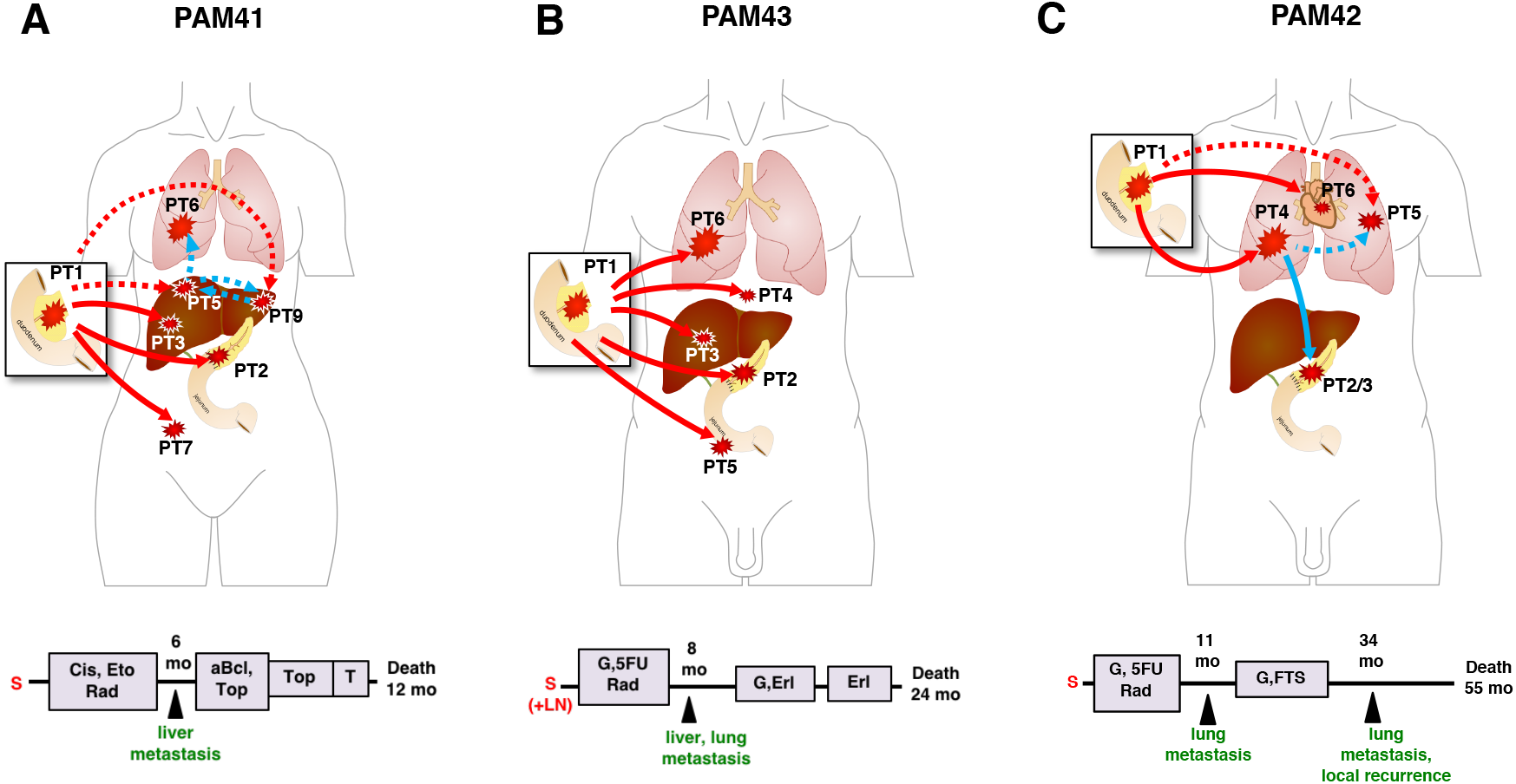
Origins of local recurrences after surgical resection in patients PAM41 (A), PAM43 (B) and PAM45 (C). The previously resected primary tumor is indicated by PT1 to the left of each patient schematic, and each patients’ clinical course is indicated below. Abbreviations are G, gemcitabine; Rad, radiation; Cis, cisplatin; Cap, capecitabine; Ox, oxaliplatin; M, Mek Inhibitor, T, Topotecan; aBcl, Bcl-2 antagonist; Erl, erlotinib, FTS, farnesylthiosalicylic acid. Solid lines indicate high confidence migration patterns and dashed lines indicate low confidence migration patterns inferred by MACHINA. Red lines reflect migration from the primary tumor and blue lines indicate migration from a site of recurrent disease. In PAM41 and PAM43 the local recurrence is seeded by the originally resected primary tumor, whereas in PAM42 it is seeded by a lung metastasis.

## Discussion

To date genomic studies of PDA have primarily relied on resections or biopsies of treatment naïve disease (14,15,42). Herein, we demonstrate that study of post-treatment samples may have value in identifying putative therapeutic vulnerabilities in recurrent disease. Of immediate clinical relevance is the finding of an increased tumor mutational burden in recurrent disease of patients treated with platinum agents, a finding that provides a rationale and context for the use of immunotherapy in the management of this disease (43). Furthermore, we also find that recurrent disease is enriched for hotspot mutations in genes associated with Mapk/Erk or PI3K/Akt signaling, some of which are potentially actionable(http://oncokb.org). Importantly, these targets are pre-existent and selected for during adjuvant treatment itself. While our sample size is sufficient for a robust gene discovery in advanced pancreatic cancer, we identified genes not commonly associated with the PDA landscape such as the nuclear exportin *XPO1*, the serine-threonine protein phosphatase *PPP6C* (24,31), or regulators of innate immunity (*PRKCI*, *TRAF3*) (34,35). These findings support the need for prospective and statistically robust efforts to sequence post-treatment PDA to better define the genes or pathways repeatedly targeted by somatic alteration. Genetic sequencing of post-treatment PDA is a critical step forward for real time molecularly informed management of recurrent disease.

Another important finding of this study is that it illustrates a taxonomy by which recurrent PDA may be stratified: monophyletic origin, polyphyletic origin or unique origin (i.e. synchronous/metachronous primaries). In all patients with bona fide recurrent PDA of mono- or polyphyletic origin, we find that systemic subclinical dissemination had already occurred at the time of surgery consistent with prior estimates (40). Questions for future investigation thus relate to methods to identify mono- versus polyphyletic recurrences in real time, and the clinical significance of this finding in the context of ongoing or planned clinical trials. Finally, while second primary carcinomas of the pancreas have been reported, our finding of a metachronous primary PDA in one of 10 otherwise unselected patients with an intrapancreatic mass post resection suggest that this phenomenon may be more common than previously appreciated (44,45). Detailed and formal prospective studies of intrapancreatic masses in patients who have undergone prior resection for PDA will be required to more firmly understand the frequency of second primary tumors and the risk factors associated with their development as has been shown for invasive carcinomas arising in intraductal papillary mucinous neoplasms of the pancreas (46).

Finally, we note that the genomic features of these patients, all of whom presented with Stage I/II disease and underwent surgery and therapy, are in stark contrast to those of untreated Stage IV PDA (47). In untreated PDA the genetic features of both the primary and metastases are remarkably uniform, and the genetic heterogeneity seen appears due exclusively to passenger mutations. By contrast, Stage I/II disease is notable for subclonal heterogeneity for driver genes as reported in other tumor types (48,49). Treatment induced genetic bottlenecks that sculpt the genomic landscape of PDA, together with intermetastatic seeding, likely contribute to subclonal and inter-metastatic heterogeneity for driver gene alterations observed in recurrent disease. Thus, context appears key in the interpretation of heterogeneity in PDA.

In summary, we identify novel genetic features of PDA in the context of recurrent disease after surgical resection and treatment with potential clinical implications for use of immunotherapy and targeted therapies in disease management. In the event that therapeutically targetable gene alterations are validated in recurrent disease, and as more targeted therapies become available, post-treatment sampling is likely to contribute to identification of the mechanisms of resistance and early identification of resistant clones.

## Methods

### Tissue Samples

Samples were generously shared by the Gastrointestinal Cancer Rapid Medical Donation Program (GICRMDP) resource at The Johns Hopkins Hospital. Sections were cut from formalin-fixed paraffin embedded (FFPE) or frozen sections available and reviewed to identify those with at least 20% neoplastic cellularity and preserved tissue quality. Samples meeting these criteria were macro-dissected from serial unstained sections before extraction of genomic DNA using DNeasy^®^ Blood & Tissue Kits for frozen samples or QIAamp DNA FFPE Tissue Kits for FFPE materials (Qiagen) following the manufacturer instructions.

### Whole Exome Sequencing

DNA quantification, library preparation and sequencing were performed in the Integrated Genomics Operation and bioinformatics analysis of somatic variants by the Bioinformatics Core at MSKCC. Libraries were created using AgilentExon_51MB_hg19_v3 as bait and sequenced on an Illumina HiSeq 2500. Whole exome sequencing resulted in a median target sequence depth of 317X (min-max) with 83% of the coding regions covered at least 100x and a calculated average tumor cellularity of 35.7% (Table S3). Paired-end sequencing data were aligned to the reference human genome (hg19) using BWA (50). Read de-duplication, base quality recalibration, and multiple sequence realignment were performed using the Picard Suite and GATK (51,52). Point mutations and small insertions and deletions were detected with MuTect and HaplotypeCaller, respectively (53). Genome-wide total and allele-specific DNA copy number was determined using the FACETS algorithm (54). For targeted sequencing of 410 cancer genes, barcoded libraries from patient-matched tumor and normal samples were captured and sequenced using methods previously reported in detail (19).

### Filtering and Annotation of Variants

For each patient, somatic variants were filtered using the following criteria: patient-matched normal coverage ≥10 reads, variant count in patient-matched normal ≤1, patient-matched normal variant frequency <0.02, tumor coverage ≥20 reads, and tumor variant allele frequency ≥0.05 in at least one tumor sample. The resulting list of somatic variants were filtered for those present in the coding regions only and subject to further bioinformatic annotation for pathogenicity and germline allele frequencies from healthy populations distributed worldwide using LiFD (55)(Table S5). MSI status was inferred from the sequencing data using a clinically validated algorithm (56), with MSI-H defined as an MSIsensor score ≥10. Copy number alterations and whole genome duplication were inferred by FACETs (54).

### Phylogenetic analysis

We applied Treeomics v1.6.0 to reconstruct the phylogenies of recurrent disease using high quality somatic variants identified by whole exome sequencing (37). Treeomics uses a Bayesian inference model to account for sequencing errors and low purity and employs Integer Linear Programming to infer a maximum likelihood tree.

### Mathematical modeling

To calculate the minimal required time a metastasis founding cell needs to grow to a detectable lesion of 1 cm3 (~109 cells) we used the smallest previously measured PDA metastasis doubling time of 27 days (57) leading to an exponential growth rate of r=0.026 per day. Assuming a PDA cell division time of 2.3 days (58), the expected minimal time for a tumor conditioned on survival takes to reach 10^9^ cells is 1.82 years (90% CI: 1.61 – 2.05 years).

### Mutational signatures

Mutational signatures were derived using the methods described by Alexandrov et al (21). To enrich for the most abundant signatures we merged those with similar putative etiologies into a single group. Signature groups present in at least 20% abundance in at least one sample were included for additional study and statistical analysis.

### Droplet Digital PCR

Absolute quantification of mutant alleles was determined using a RainDrop Plus Digital PCR system according to the manufacturer instructions. Predesigned or custom designed TaqMan assays were obtained for variants of interest (Thermo Fisher). Approximately 75 ng of gDNA were used per reaction in a 25 ul volume. Each reaction contained 5 million droplets at a target inclusion rate of 10% (500,000 target molecules).

### Migration Pattern Inferences

PyClone (59) and MACHINA (60) were used to infer seeding patterns associated with metastasis and local recurrence. To alleviate long run times associated with the high sample number context we applied specific filters to focus on the most informative loci. These were a) the locus was sequenced to a depth of at least 60 in at least one sample; b) the locus had a copy number profile consistent with a relatively simple genomic history in all samples (the combination of major allele A and minor allele B at the locus was required to be one of AB, AAB, AAAB, or AABB in each sample, though not necessarily the same across samples) and c) each locus was required to be sequenced to nonzero depth in all remaining samples. Samples that did not contain more than 20 variant loci after applying this filter were excluded. In patient PAM41, a large cluster of highly related liver metastases (PT4, 8, 9, 10, 11) were simplified by including only sample PT9 to improve runtime. The resulting mutations and associated major and minor copy numbers were clustered with PyClone using default settings. The PyClone consensus cluster files were used to enumerate evolutionary relationships, then the combination of PyClone cluster frequency estimates and enumerated trees together were used to search for the most parsimonious migration patterns consistent with each tree topology. The resulting solution with the lowest overall migration number, and then the lowest comigration number, were determined. No other constraints were applied to the migration plots.

### Statistics

Descriptive data are represented as a mean and standard deviation unless otherwise mentioned. Parametric distributions were compared by a Student’s T-test whereas non-parametric distributions were compared by a Mann-Whitney U test. Frequency data were compared using a X^2^ test. All comparisons were two-sided. Survival curves were plotted according to the methods of Kaplan and Meier and compared using a log-rank test.

## Acknowledgements

Supported by NIH/NCI grants R01 CA179991 and R35 CA220508 to C.I.D., 2T32 CA160001-06 to A.M.M, the Daiichi-Sankyo Foundation of Life Science Fellowship to A.H, the Mochida Memorial Foundation for Medical and Pharmaceutical Research Fellowship to A.H, R00 CA22999102 to J.G.R., and NIH/NCI P50 CA62924. This work was funded in part by the Marie-Josée and Henry R. Kravis Center for Molecular Oncology and the National Cancer Institute Cancer Center Core Grant No. P30-CA008748.

