## Supplementary material for "Evolutionary Origins of Recurrent Pancreatic Cancer": Figures S1 and S2

by

Hitomi Sakamoto, Marc A. Attiyeh, Jeffrey M. Gerold, Alvin P. Makohon-Moore, Akimasa Hayashi, Jungeui Hong, Rajya Kappagantula, Lance Zhang, Jerry Melchor, Johannes G. Reiter, Alexander Heyde, Craig M. Bielski, Alexander Penson, Mithat Gönen, Debyani Chakravarti, Eileen M. O'Reilly, Laura D. Wood, Ralph H. Hruban, Martin A. Nowak, Nicholas D. Socci, Barry S. Taylor, Christine A. Iacobuzio-Donahue*

This file contains:

**Supplemental Figure 1**

**Supplemental Figure 2**

*Supplemental Tables 1-7 are in a separate file.*

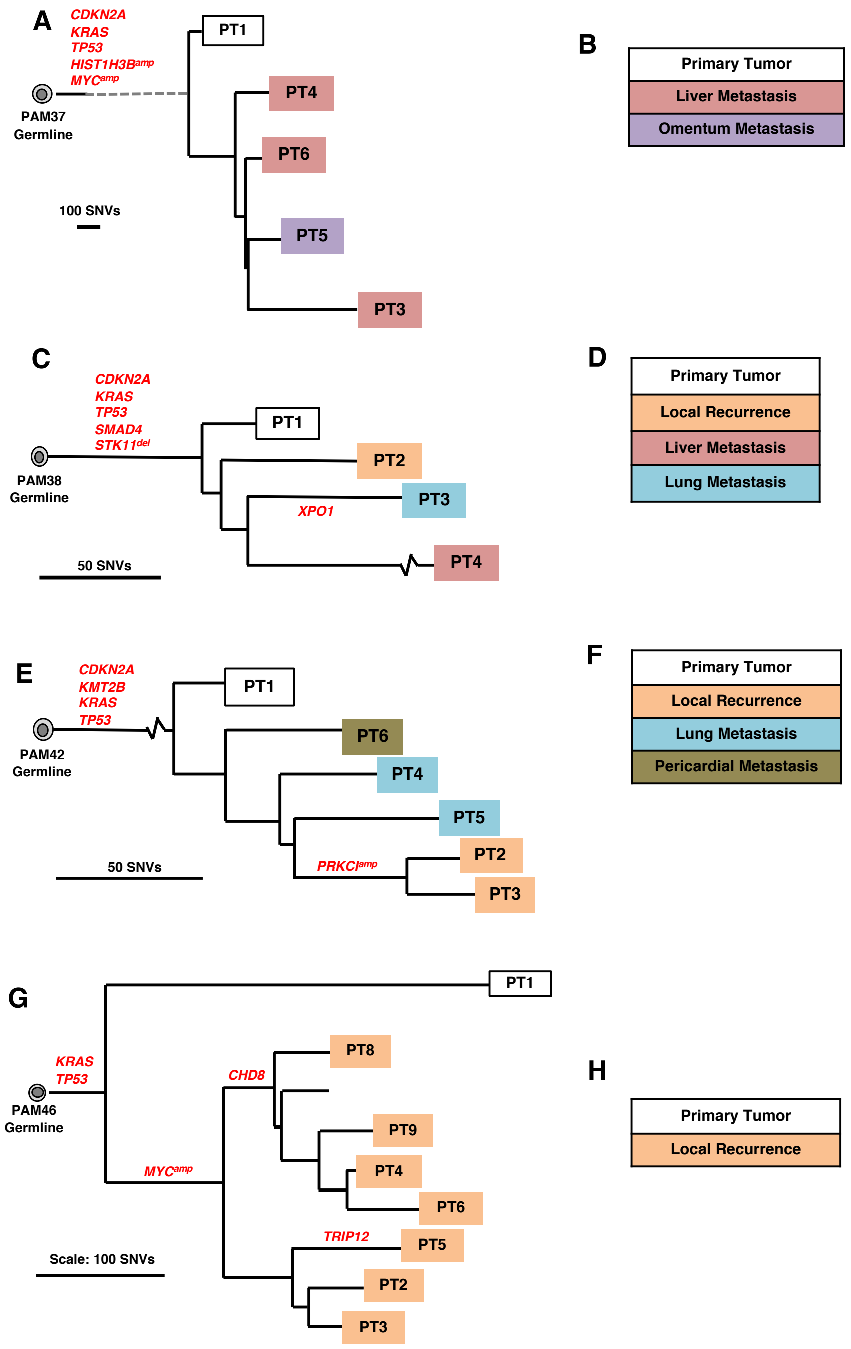


**Supplemental Figure 1: Monophyletic origins of recurrent pancreatic cancer**. Shown are the phylogenetic analyses of the relationships of the primary tumor to the local recurrence and liver metastases in patients PAM37 (**A**), PAM38 (**C**), PAM42 (**E**) and PAM46 (**G**). Color code of sample origins for each patient are shown in **B, D, F** and **H**. In all four patients the recurrent disease has a common ancestor and the primary tumor (PT1) is the outgroup in the tree. Subclonal driver gene alterations are also seen in patients PAM38 (*XPO1*), PAM42 *(PRKCI*) and PAM46 (MYC, CHD8, TRIP12).

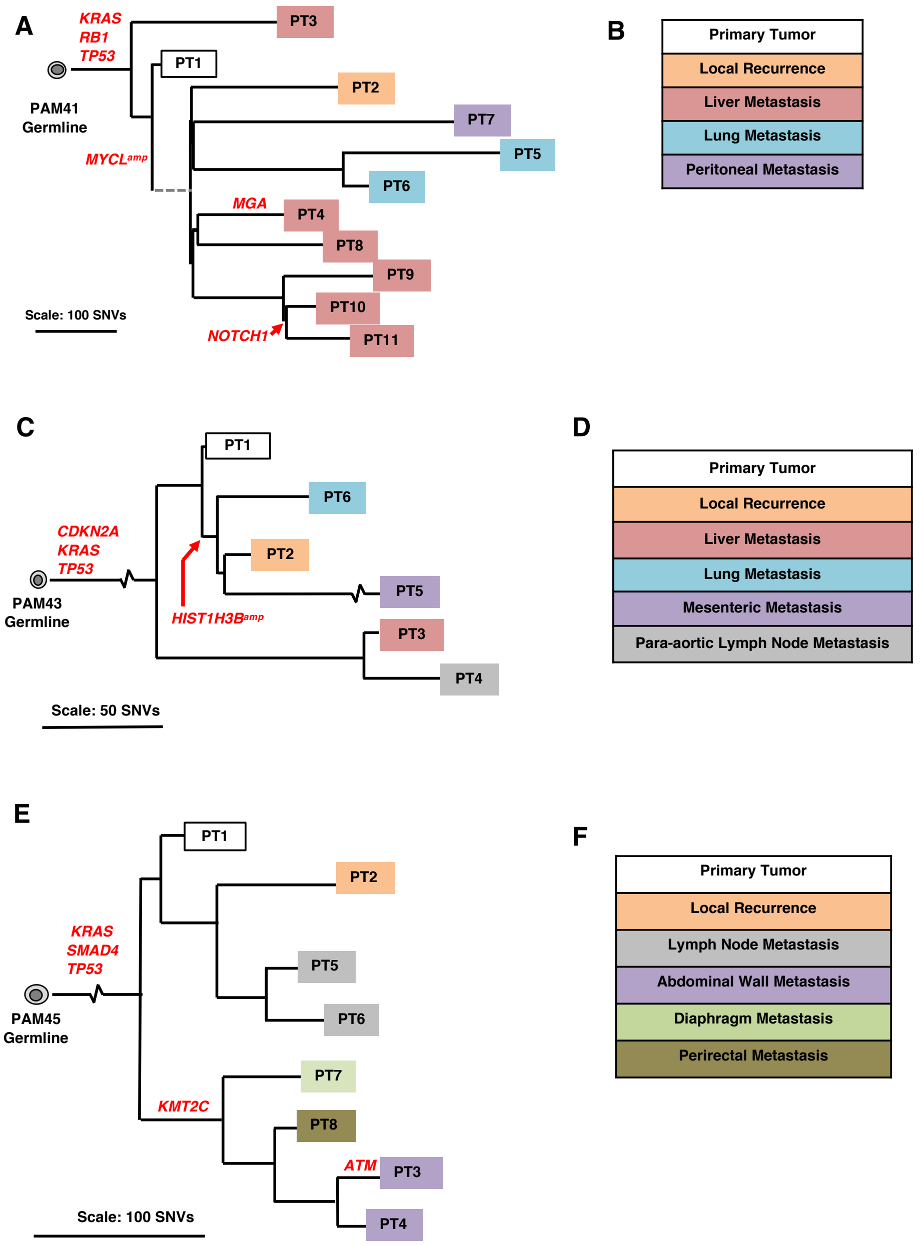


**Supplemental Figure 2: Polyphyletic origins of recurrent pancreatic cancer**. Shown are the phylogenetic analyses of the relationships of the primary tumor to the local recurrence and distant metastases in patients PAM41 (**A**), PAM43 (**C**), and PAM45 (**E**). Color code of sample origins for each patient are shown in **B, D,** and **F**. In all three patients there is no common ancestor for the recurrent disease. Subclonal driver gene alterations are also seen in all three patients.
